## Supporting Information for "Sld3CBD–Cdc45 structural insights into Cdc45 recruitment for CMG complex formation on DNA replication"

**Supplementary Figure 1**

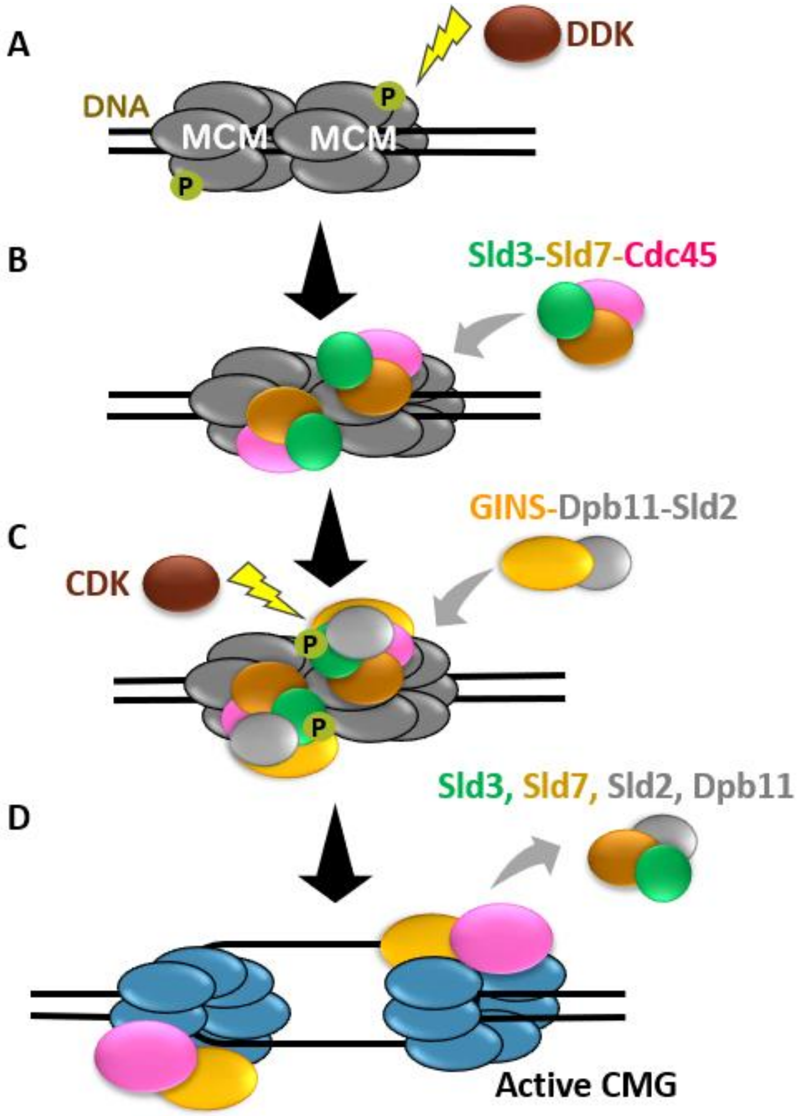

**Supplementary Figure 1. Formation of the CMG complex after MCM DH bound to the replication initiation site**

(A) MCM DH binds to the replication origin and then is phosphorylated by DDK.
(B) Sld3 and Sld7 recruit Cdc45 to the MCM DH complex. (C) After CDK phosphorylates Sld3, Dbp11 and Sld2 recruit GINS to MCM DH to form the active helicase CMG complex.
(D) Finally, factors other than CMG (Cdc45–MCM–GINS) are dissociated, and dsDNA is unwound to ssDNA in preparation for initiating replication.

**Supplementary Figure 2**

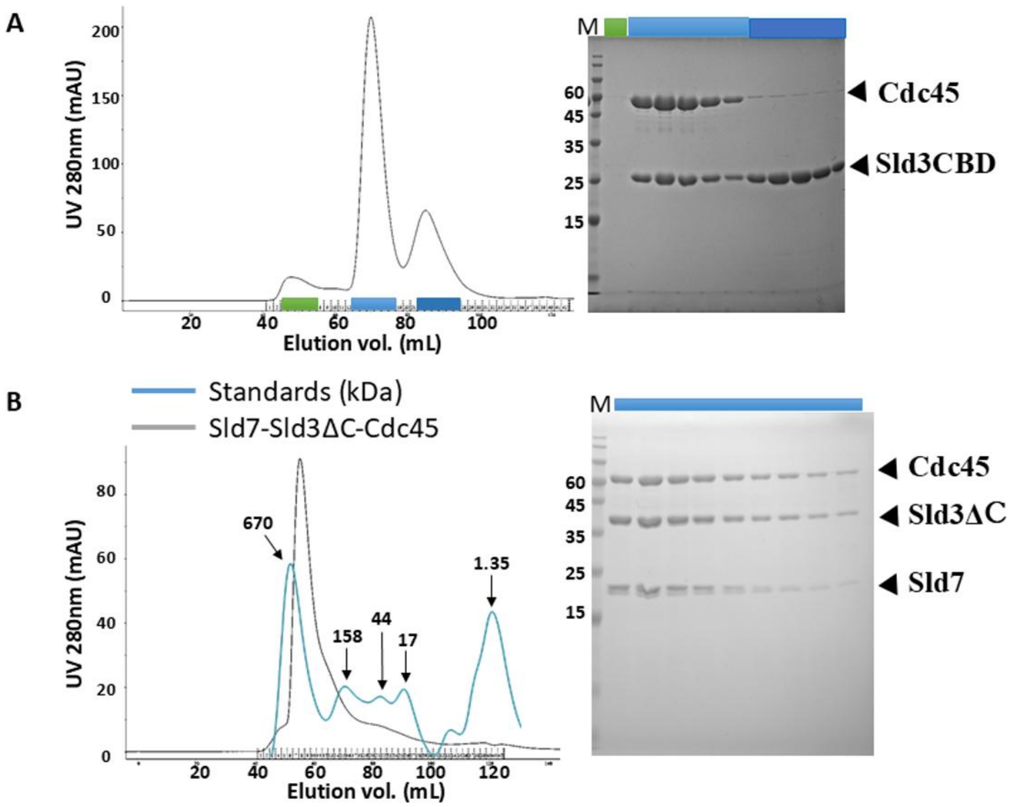

| | Thyroglobu lin (bovine) | g-globulin (bovine) | Ovalbumin (chicken) | Myoglobin (horse) | Vitamin B12 | Sld7-Sld3ΔC-Cdc45 (Data fitting) |
|---|---|---|---|---|---|---|
| M.W. (kDa) | 670 | 158 | 44 | 17 | 1.35 | 429 |
| Elu. (mL) | 51.7 | 70.1 | 82.8 | 90.9 | 131.2 | 55.8 |

**Supplementary Figure 2. Purification of the Sld3CBD–Cdc45 and Sdl7–Sld3ΔC–Cdc45 complexes**

Size-exclusion chromatography (SEC) of ScSld3CBD–Cdc45 (A) and KmSld7–Sld3ΔC–Cdc45 (B). SDS-PAGE was performed to examine the purity of each sample in the SEC plots. Lane M shows the molecular weight markers labelled in kDa. At the upper left, we collected the principal peak in the SEC as Sld3CBD–Cdc45, which was used in other experiments. According to the bottom-right image, we collected the first half of the marked peak in SEC (B left) as purified Sld7–Sld3ΔC–Cdc45. A standards kit was measured using Superdex 200 16/60 column to check the elution volume shift of Sld7–Sld3ΔC–Cdc45. The molecular weight at the peak elution position of Sld7-Sld3ΔC-Cdc45 was estimated to be 429 kDa.

**Supplementary Figure 3**

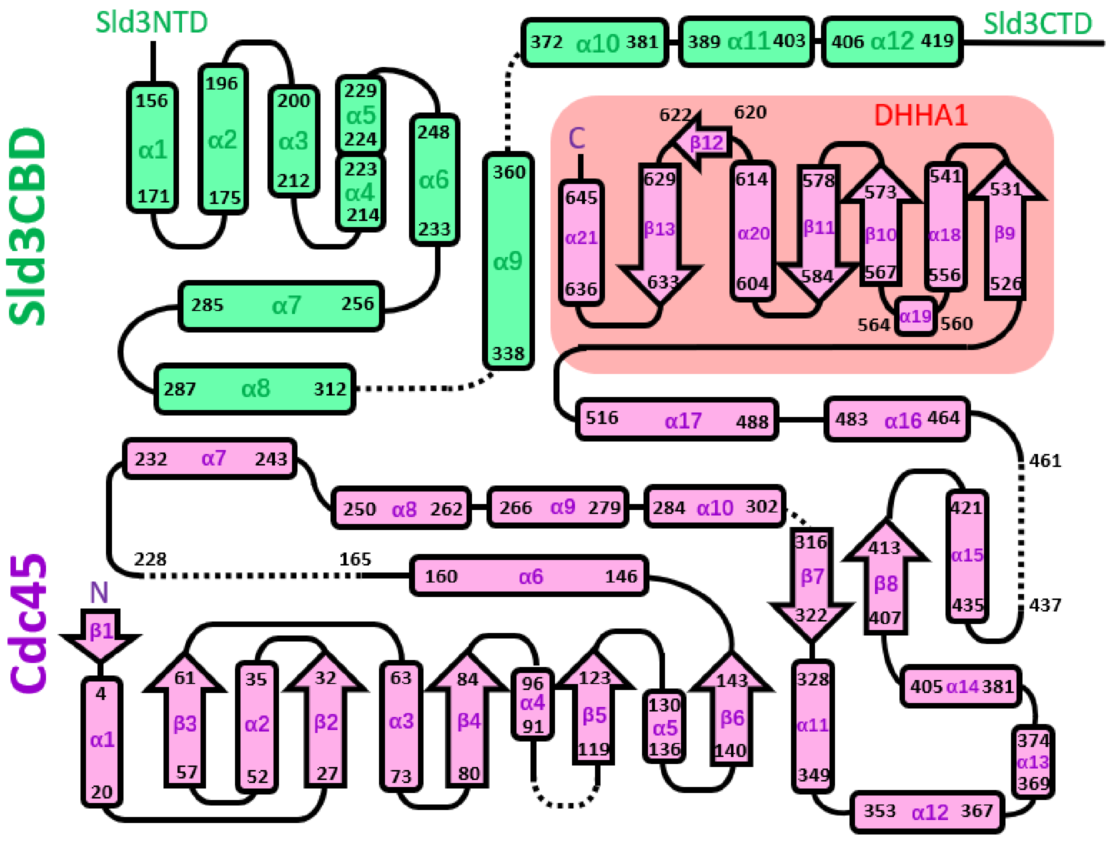

**Supplementary Figure 3. The topology diagram of Sld3CBD-Cdc45**

The topology diagram of Sld3CBD and Cdc45 in the Sld3CBD-Cdc45 complex structure. α helix and β sheet were shown by the square and the arrow, respectively. The black line and dotted line represent the loop and disorder regions, respectively.

### Supplementary Figure 4

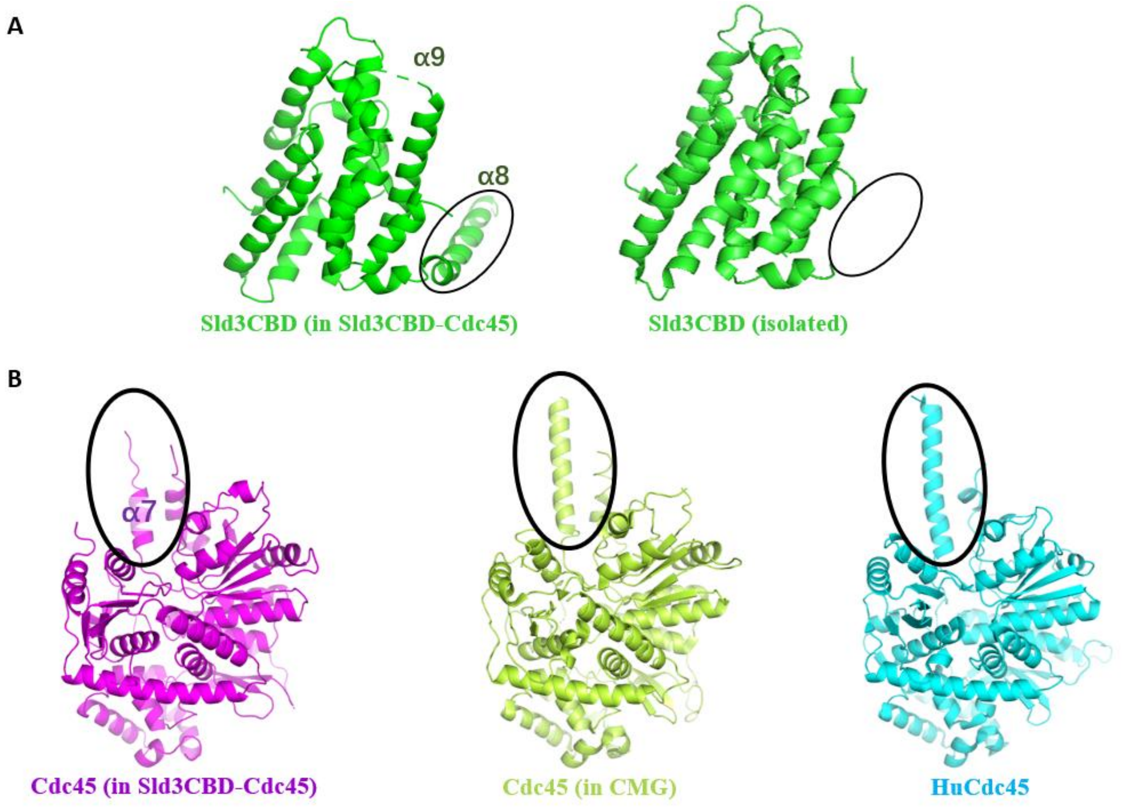

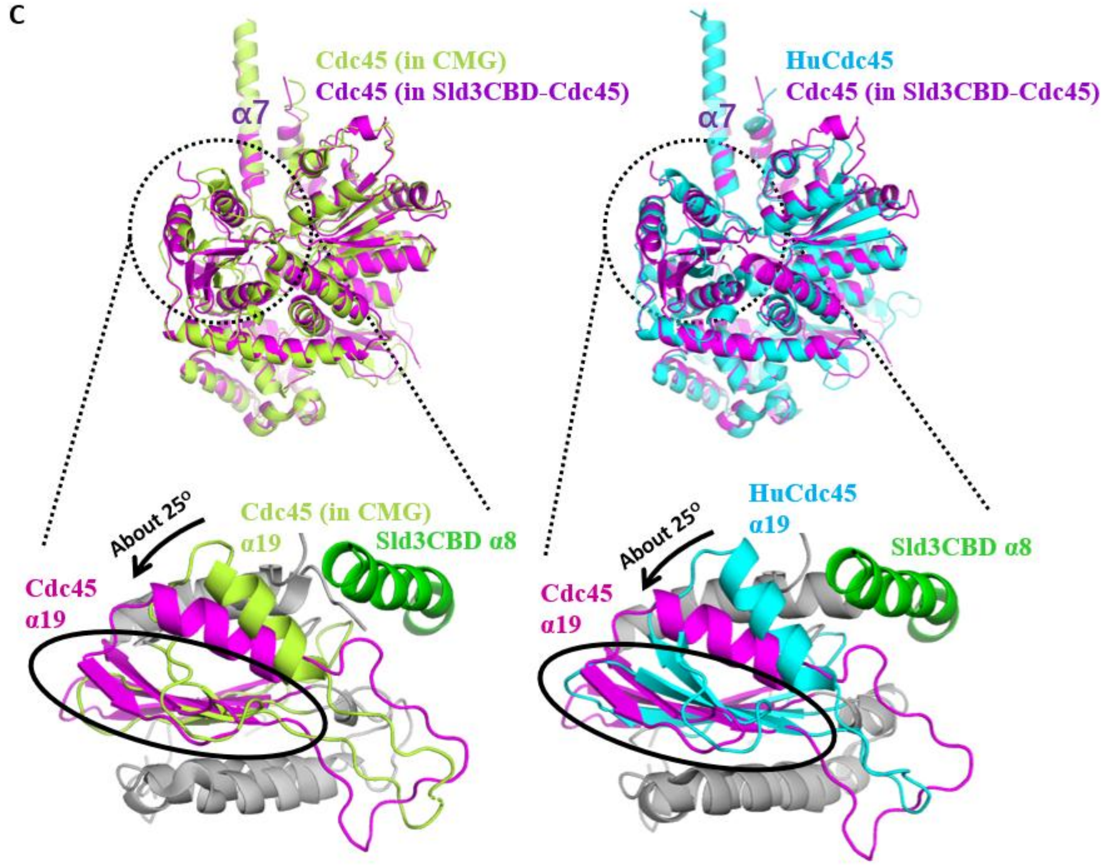

**Supplementary Figure 4. Structural Comparison of Sld3CBD and Cdc45**

(A) Sld3CBD in the Sld3CBD–Cdc45 complex (left) and the isolated structure of Sld3CBD (3WI3) (right). The black circle indicates a long helix α8CTP that was only visible in the Sld3CBD–Cdc45 complex with an average B factor of 45 $A^2$ for the main chain. (B) Cdc45 in Sld3CBD–Cdc45 (left; magenta), CMG (3JC6) (center; yellow-green), and isolated HuCdc45 structure (5DGO) (right; cyan). In Sld3CBD–Cdc45, the black circle indicates a long helix that was partly disordered. (C) Superposition of Cdc45 in Sld3CBD–Cdc45 by aligning Cdc45NTD (~K517) to Cdc45 in the CMG complex and huCdc45, respectively (upper panel); colors are the same as those in (B). The black dotted circles indicate conformationally changed DHHA1 domains (magnified below the images). Significant changes in the α19 and downstream β-sheets in the DHHA1 domain are highlighted by the black arrow and circle, respectively. The parts with no significant conformational changes are colored gray, and the colors of the other parts are the same as those in (B).

### Supplementary Figure 5

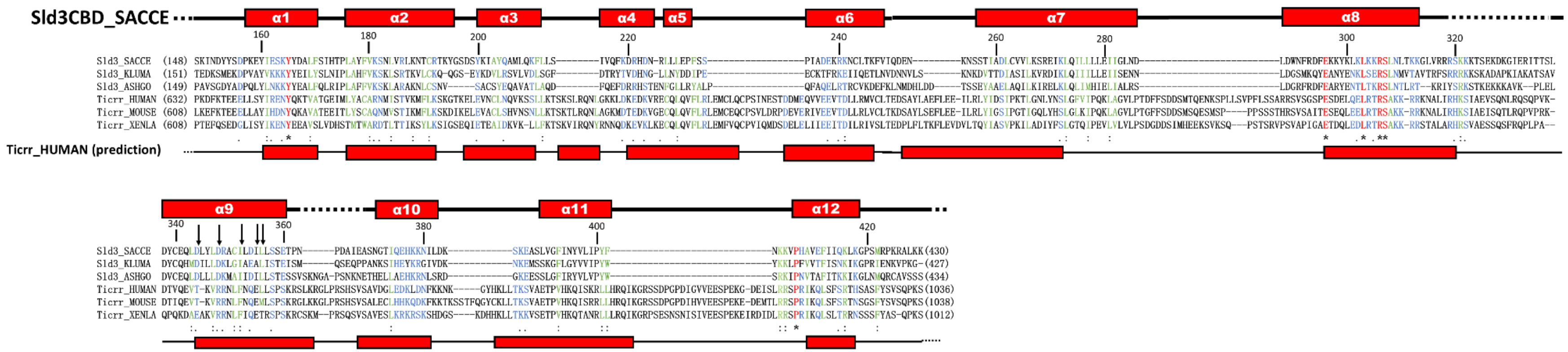

**Supplementary Figure 5. Sequence alignment of Sld3/Treslin domain with structural elements**

Sequence alignment of the Sld3/Treslin domain (Cdc45-binding domain: CBD) with structural elements from fungal Sld3 (*S. cerevisiae*, *K. marxianus,* and *Ashbya gossypii*) and vertebrate Treslin/Ticrr (*Homo sapiens*, *Mus musculus,* and *Xenopus laevis*). The sequences are as follows: Sld3_SACCE, *S. cerevisiae*; Sld3_KLUMA, *K. marxianus*; Sld3_ASHGO, *Ashbya gossypii*; Ticrr_HUMAN, *Homo sapiens*; Ticrr_MOUSE, *Mus musculus*; and Ticrr_XENLA, *Xenopus laevis*.

CLUSTAL W (https://www.genome.jp/tools-bin/clustalw) was used to create an initial alignment, which was modified based on the 3D structure. Structural elements of TICRR_HUMAN were predicted using PSIPRED 4.0 (http://bioinf.cs.ucl.ac.uk/psipred/). * conserved sequence; **:** and **.** conserved change. Amino acids are marked and colored red, green, and blue. The secondary structures of Sld3CBD (in Sld3CBD–Cdc45) and predicted TICRR_HUMAN are shown above and below the alignment, respectively. Dashed lines in the secondary structure indicate disordered regions. The residue numbers for SLD3_SACCE are indicated above the alignment. The numbers in parentheses beside the sequences indicate the number of residues in each protein. The mutation sites used in this study are indicated by black arrows.

### Supplementary Figure 6

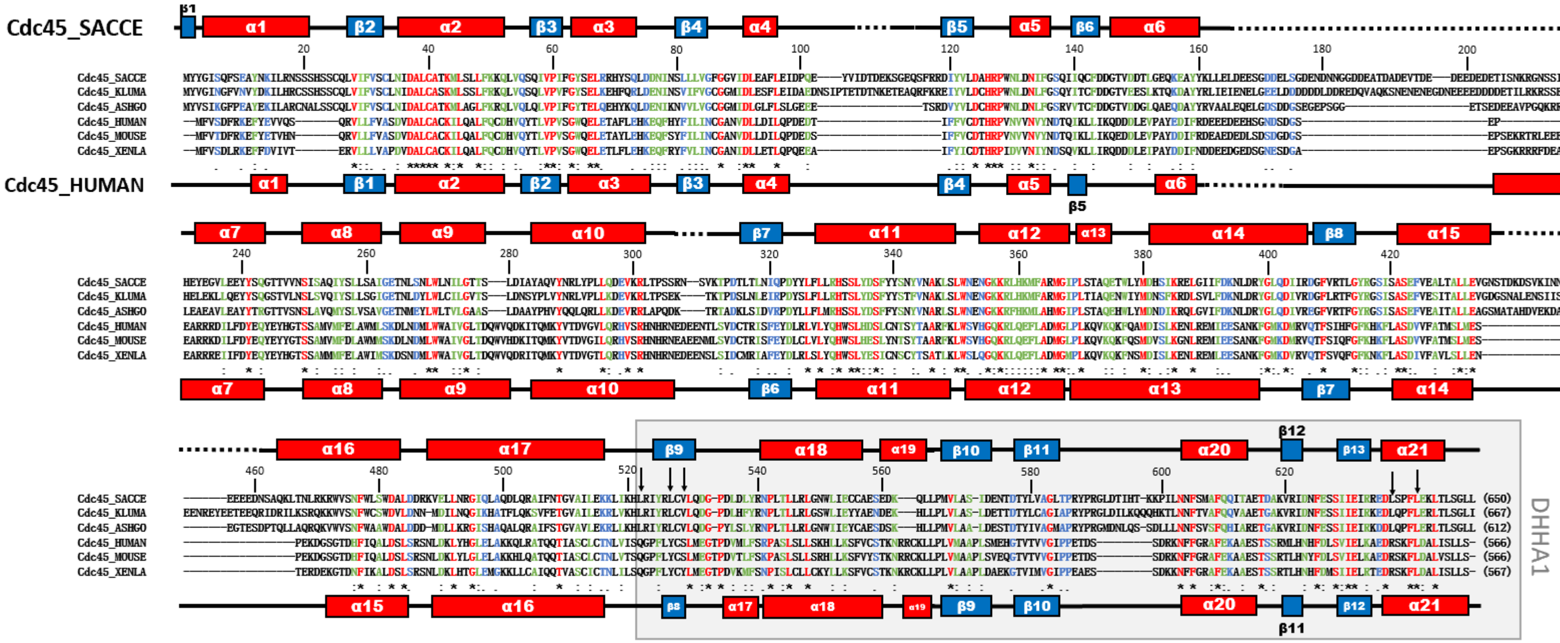

**Supplementary Figure 6. Sequence alignment of Cdc45s with structural elements**

Sequence alignment of Cdc45s with structural elements from fungi (*S. cerevisiae*, *K. marxianus,* and *Ashbya gossypii*) and vertebrates (*Homo sapiens*, *Mus musculus,* and *Xenopus laevis*). The sequences are indicated as Cdc45_SACCE, *S. cerevisiae*; Cdc45_KLUMA, *K. marxianus*; Cdc45_ASHGO, *Ashbya gossypii*; Cdc45_HUMAN, *Homo sapiens*; Cdc45_MOUSE, *Mus musculus*; and Cdc45_XENLA, *Xenopus laevis*.

CLUSTAL W (https://www.genome.jp/tools-bin/clustalw) was used to create an initial alignment, which was modified based on the 3D structure. ***** conserved sequence; **:** and **.** conserved change. Amino acids are marked and colored red, green, and blue. Secondary structures of Cdc45 in Sld3CBD–Cdc45 and HuCdc45 (PDBID: 5DGO) are indicated above and below the alignment, respectively. Dashed lines in the secondary structure indicate disordered regions. The residue numbers for Cdc45_SACCE are shown above the alignment. The numbers in parentheses beside the sequences indicate the number of residues in each protein. Black arrows indicate the mutation sites used in this study. A black frame shows the domain DHHA1.

**Supplementary Figure 7**

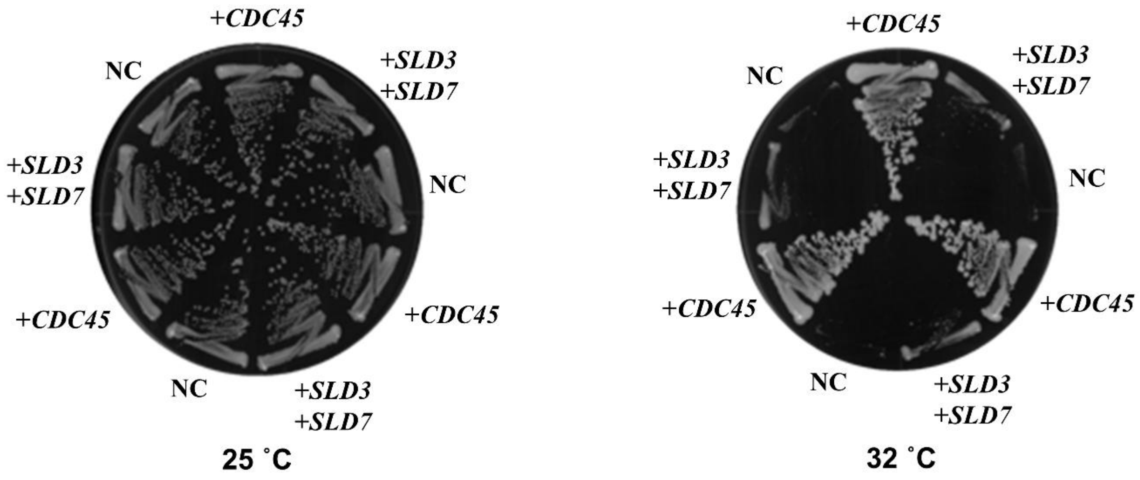

**Supplementary Figure 7. *In vivo* mutation analysis of Cdc45 using mutant cells**

*SLD3* on the high-copy YEplac195 plasmid and *SLD7* on the high-copy YEplac112 plasmid were introduced into mutant cells bearing a mutation that replaced Cdc45 Ser242 with proline on the Sld3 binding surface (Cdc45S242P). The transformants were streaked onto yeast extract–peptone–dextrose plates and individually incubated for 3 days at 25°C and 32°C. Blank plasmids YEplac195 and YEplac122 were used as negative controls (NC). *CDC45* was introduced into the YEplac195 plasmid as a positive control. Cell growth was suppressed by Cdc45S242P mutation at 32°C.

**Supplementary Figure 8**

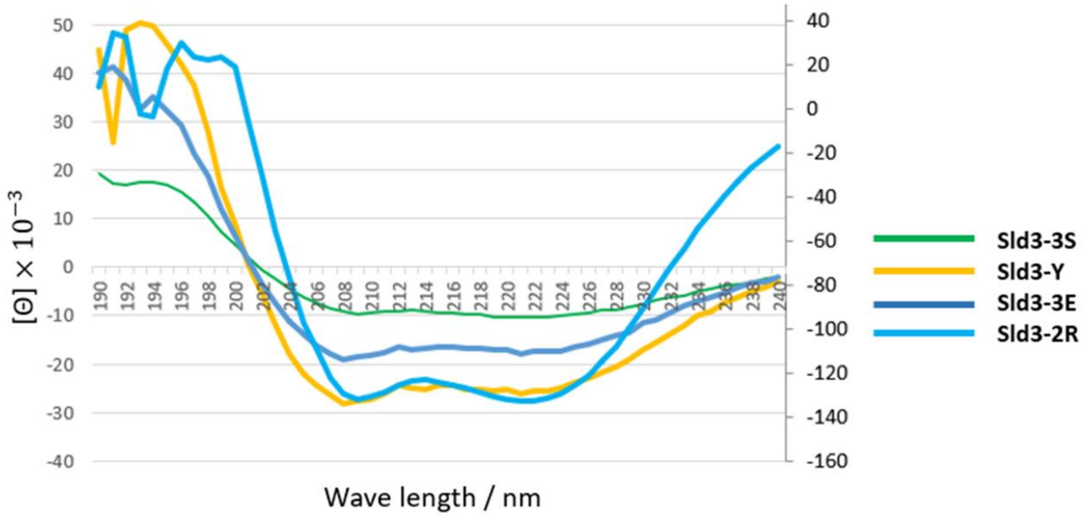

| | α helix | β strand |
|---|---|---|
| Sld3CBD(-Cdc45) | 78.8% (16.3% disordered) | 0% |
| Sld3-3S | 81.79% | 1.44% |
| Sld3-Y | 80.99% | 1.83% |
| Sld3-3E | 81.29% | 1.80% |
| Sld3-2R | 78.03% | 1.92% |

**Supplementary Figure 8. SDS-PAGE analysis and circular dichroism spectra of Sld3 mutants**

Structural elements of Sld3-3S, Sld3-3E, Sld3-2R, and Sld3-Y (Sld3-3S: I352S/I355S/L356S, Sld3-3E: 352E/I355E/L356E, Sld3-2R: D344R/D348R, and Sld3-Y: I352Y) were analyzed through circular dichroism. While the mutants of Sld3CBD existed alone, we prepared WT Sld3CBD in a complex with Cdc45 and calculated the elements of secondary structure from the crystal structure of Sld3CBD–Cdc45. All variants appeared to maintain the same structural elements as wild-type Sld3CBD–Cdc45, as indicated in the table. The concentration of samples was controlled to the same level for CD measurement.

**Supplementary Figure 9**

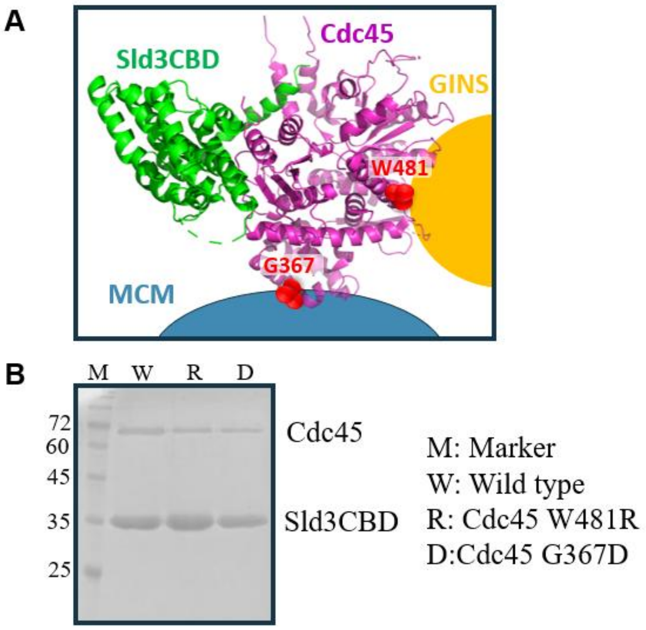

**Supplementary Figure 9 Mutation analysis of Cdc45**

(A) Binding site of Cdc45 to Sld3CBD, GINS and MCM. Two Cdc45 sites involved in binding to MCM (Cdc45 G367) and GINS (Cdc45 W481) are colored in red. (B) *In vitro* binding analysis was checked using SDS-PAGE after Ni-affinity chromatography extraction of co-overexpressed Sld3CBD with each Cdc45 mutant. Sld3CBD with His-tag bound to the column. The labels M, W, R and D are explained on the right.

**Supplementary Figure 10**

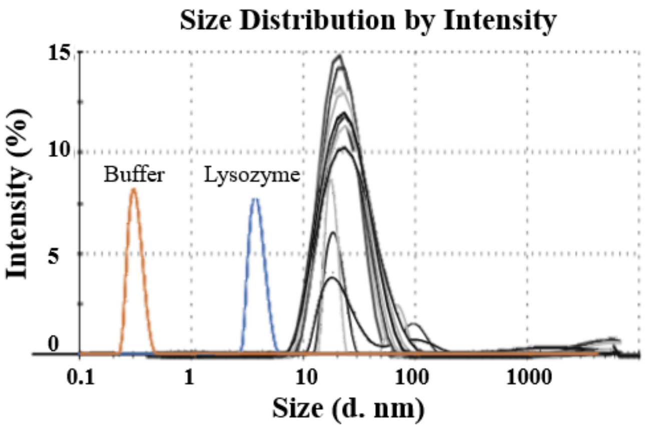

| | | |
|---|---|---|
| Buffer control: | 0.30 (d. nm) | |
| Standard (Lysozyme) : | 4.11 (d. nm) | |
| | Size (d. nm) | % Intensity |
| Sample 1 | 23.59 | 81.4 |
| Sample 2 | 23.42 | 95.6 |
| Sample 3 | 23.10 | 100 |
| Sample 4 | 22.60 | 95.1 |
| Average | 23.18±0.22 | |

**Supplementary Figure 10. Dynamic light scattering (DLS) of Sld7–Sld3ΔC–Cdc45**
Four samples of the Sld7–Sld3ΔC–Cdc45 complex were measured by DLS. Each sample was overexpressed independently and purified through size-exclusion chromatography. For data analysis, each sample was measured three times. The particle size was estimated to be 232 Å in average of peak size from four samples. The pure buffer was measured as a background control and 10 μM lysozyme was measured as a standard control, show by orange and blue curves, respectively

**Supplementary Figure 11**

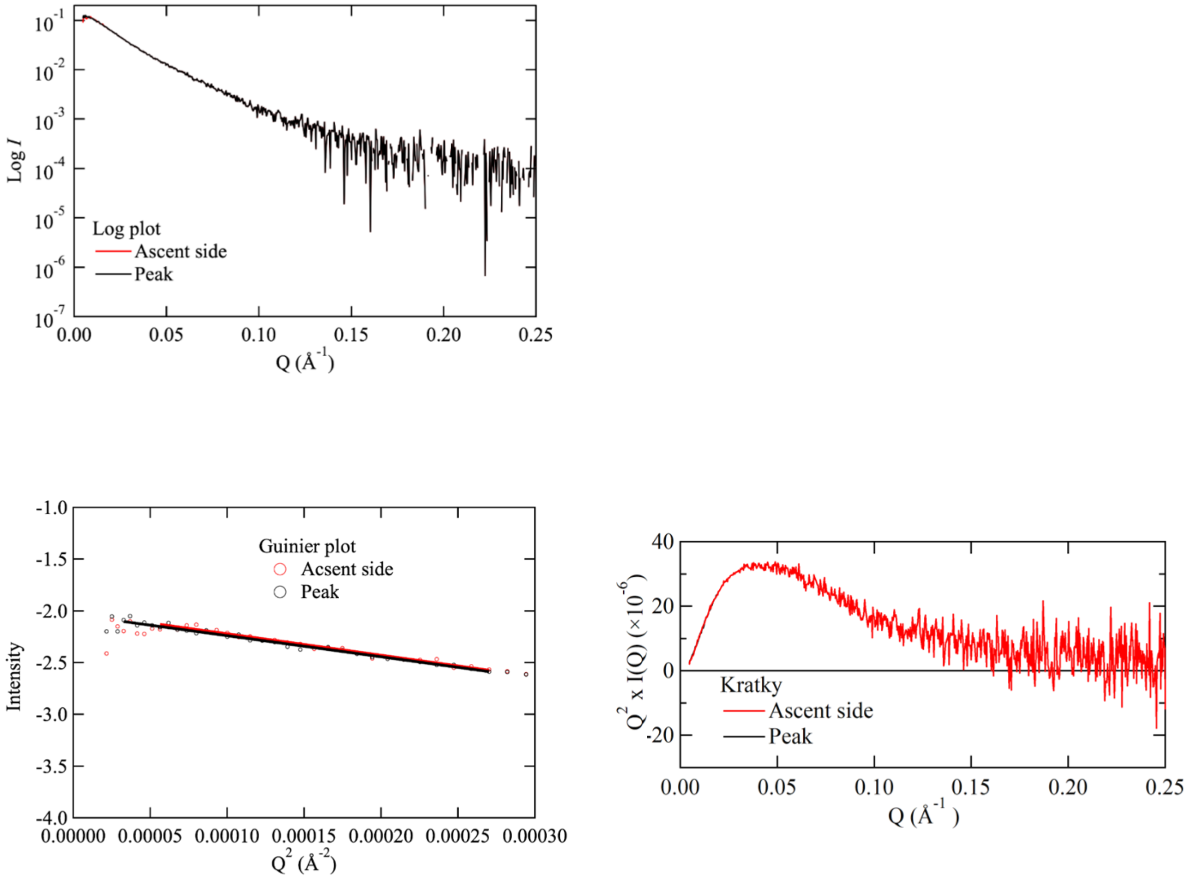

**Supplementary Figure 11. SEC-SAXS analysis of Sld7–Sld3ΔC–Cdc45**

SEC-SAXS measurements of Sld7–Sld3ΔC–Cdc45 complex were performed using a beam line BL-10C at the Photon Factory (Tsukuba, Japan). SEC-SAXS data were collected under camera length 2 M, wavelength 1.5Å and 20℃ with a program Seral Analyzer (upper panel) [1]. A program SAngler was used to analyze the SEC-SAXS data [2]. The Guinier plot (left) and Kratky plot (right) are shown in the lower panel. Rg and Dmax were estimated around 85 and 345 Å, respectively. Acsent side: the SAXS data collated from the left side of the SEC-plot peak. Peak: the SAXS data collected from a peak point of the SEC plot.

### Supplementary Figure 12

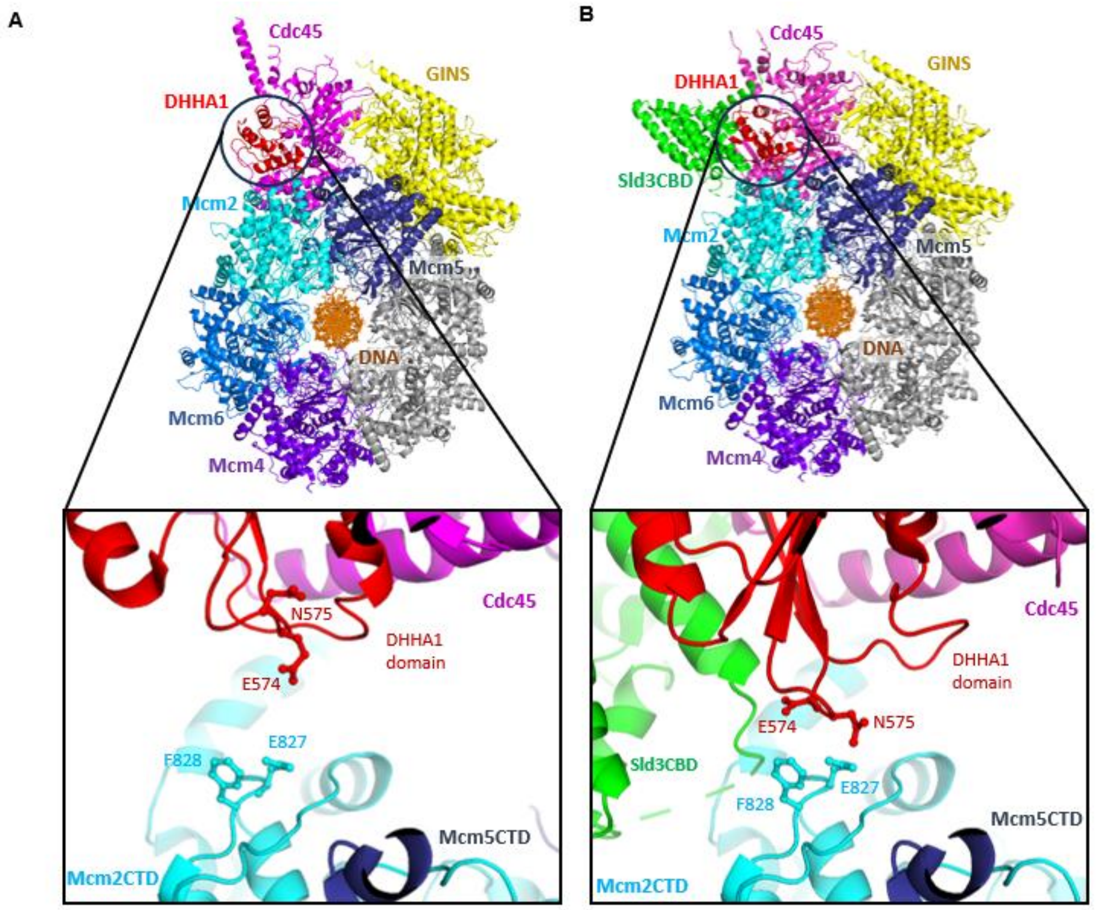

**Supplementary Figure 12. DHHA1 domains of Cdc45s**

The DHHA1 domains of Cdc45 in the CMG complex (PDB ID: 3JC6) (A) and the SCMG–dsDNA model (B), are shown. DHHA1s are colored in red and around the black circles. Labelled Mcm2, 5, 4, and 6 subunits are colored in cyan, blue, marine, and light blue, respectively. Subunits Mcm3 and Mcm7 are colored in gray. Green and pink were used to indicate Sld3CBD and Cdc45, respectively. GINS is shown in yellow and dsDNA is presented as a dark orange stick. The contact area between DHHA1 and Mcm2 is magnified in the bottom panel of the figure. The black dotted circles mark the contact between DHHA1 and Mcm2CTD in CMG complex (A) and SCMG–dsDNA model (B), respectively. The predicted closed residues between Cdc45 and Mcm2 in the SCMG–dsDNA model are labelled.

### Supplementary Figure 13

A

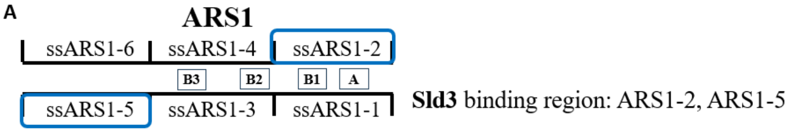

ssARS1-1 **5′- TTACATCTTGTTATTTTACAGATTTTATGTTTAGATCTTTTATGCTTGCTTTTCAAAAGGCCTGCAGGCAAGTGCACAAA-3′**
ssARS1-2 **3′-AATGTAGAACAATAAAATGTCTAAAATACAAATCTAGAAAATACGAACGAAAAGTTTTCCGGACGTCCGTTCACGTGTTT-5′**

ssARS1-3 5′- CAATACTTAAATAAATACTACTCAGTAATAACCTATTTCTTAGCATTTTTGACGAAATTTGCTATTTTGTTAGAGTCTTT-3′
ssARS1-4 **3′- GTTATGAATTTATTTATGATGAGTCATTATTGGATAAAGAATCGTAAAAACTGCTTTAAACGATAAAACAATCTCAGAAA-5′**

ssARS1-5 5′- TACACCATTTGTCTCCACACCTCCGCTTACATCAACACCAATAACGCCATTTAATCTAAGCGCATCACCAACATTTTCTG-3′
ssARS1-6 **3′-ATGTGGTAAACAGAGGTGTGGAGGCGAATGTAGTTGTGGTTATTGCGGTAAATTAGATTCGCGTAGTGGTTGTAAAAGAC-5′**

B

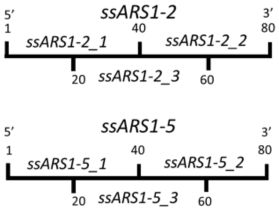

| **ssDNA** | **Sequences** |
|---|---|
| **ssARS1-2_1** | TTTGTGCACTTGCCTGCAGGCCTTTTGAAAAGCAAGCATA |
| **ssARS1-2_2** | AAAGATCTAAACATAAAATCTGTAAAATAACAAGATGTAA |
| **ssARS1-2_3** | CCTTTTGAAAAGCAAGCATAAAAGATCTAAACATAAAATC |
| **ssARS1_5_1** | TACACCATTTGTCTCCACACCTCCGCTTACATCAACACCA |
| **ssARS1-5_2** | ATAACGCCATTTAATCTAAGCGCATCACCAACATTTTCTG |
| **ssARS1-5_3** | CTCCGCTTACATCAACACCAATAACGCCATTTAATCTAAG |

**Supplementary Figure 13. Sequence of ssARS1 fragments**

(A) Fragments of ssARS1-1 through ssARS1-6 used in this study. Each ssARS1 fragment contains 80 bs. Fragments of odd and even numbers are complementary strands. The important elements A (ARS consensus sequence), B1, B2, and B3 for unwinding are marked [28]. The fragments in the blue square are Sld3 binding parts of ssARS1. (B) The fragments of Sld3 binding parts (blue square parts in (A)) of ssARS1-2 and ssARS1-5. Each fragment of ssARS1-2 and ssARS1-5 is separated into 40 bs lengths: ssARS1-2-1 to ssARS1-2-3 and ssARS1-5-1 to ssARS1-5-3.

**Supplementary Figure 14**

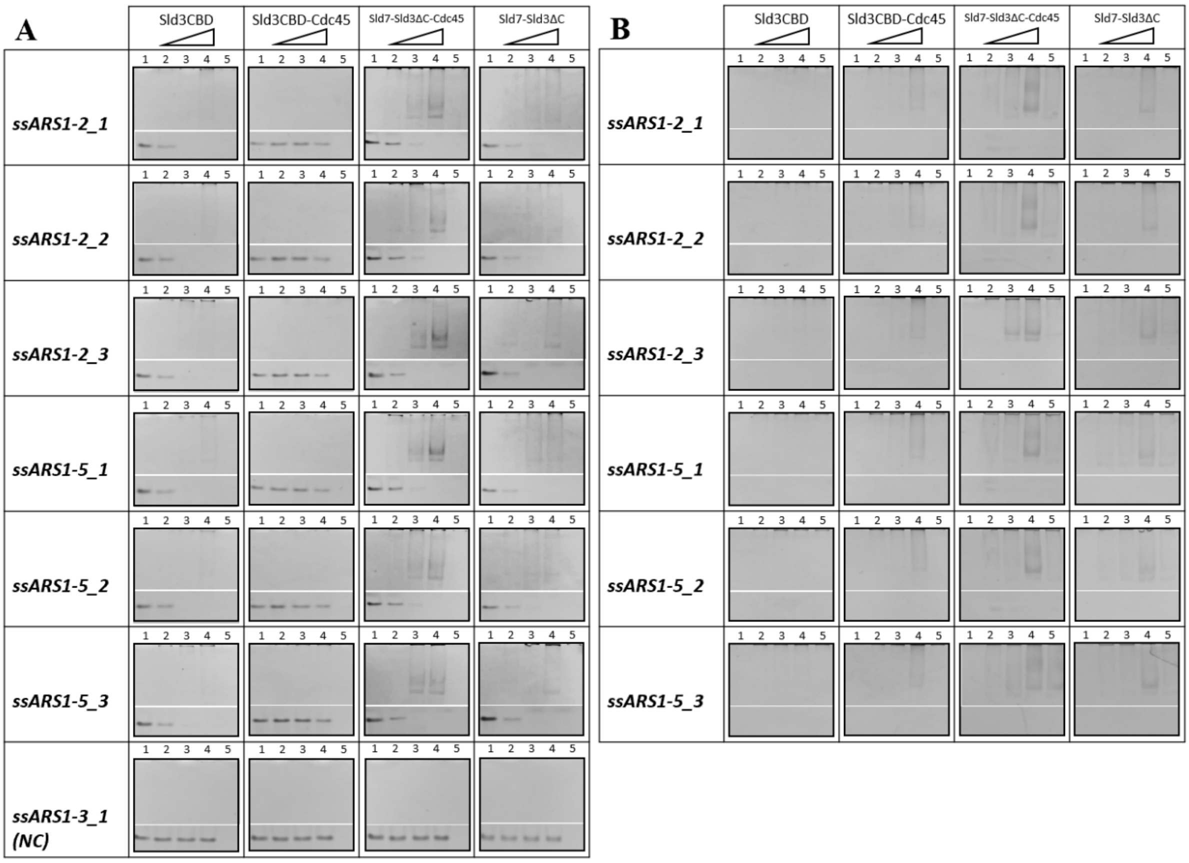

**Supplementary Figure 14. Electrophoresis mobility shift assay (EMSA) of ssDNA binding to Sld3 and its complexes with Sld7 and Cdc45**

(A) ssDNAs were visualized using Fast Blast DNA stain on polyacrylamide gels. In the presence of ssDNA fragments, Sld3CBD, Sld3CBD–Cdc45, Sld7–Sld3ΔC–Cdc45 and Sld7–Sld3ΔC were incubated with molecular mass-related concentrations. The molecular ratio of ssDNA to protein in lanes 1, 2, 3, 4, and 5 was 1:0, 1:0.5, 1:1, 1:2, and 0:1, respectively. The controls for ssDNA and protein are lanes 1 and 5, respectively. The negative controls for no binding with ssDNA (ssARS 1-3_1, NC) of each sample are shown at the bottom. The reduce or disappearance of the ssDNA band in lanes 2-4 indicates that the protein (Sld3CBD, Sld7–Sld3ΔC and Sld7–Sld3ΔC–Cdc45) binds to ssDNA with high affinity. The smeared bands appear in high molecular weight regions of lanes 2-4, when mixed with Sld7–Sld3ΔC–Cdc45 or Sld7–Sld3ΔC, whereas no bands appeared in the NC (ssARS1-3_1). The positions of smeared ssDNA bonds correspond to those of protein in the protein-stain pages, indicating that ssARS1 were complexed with proteins.

(B) The proteins were visualized using Coomassie brilliant blue on gels. The EMSA experiments were conducted concurrently under equivalent conditions to (A). The smeared bands in the high molecular weight parts of lanes 2–4 of Sld3CBD–Cdc45, Sld7–Sld3ΔC–Cdc45, and Sld7–Sld3ΔC are shown more clearly when mixed with ssDNA. Such enhanced discernibility indicates that these proteins easily enter the gel with ssDNA, even though Sld3CBD–Cdc45 binds ssDNA weakly. Sld3CBD could not enter the gel, even when bound to ssDNA, because the pI values exceeded the pH of the running buffer (pH = 8.3).

Due to limitations in protein overexpression, we utilized Sld7–Sld3ΔC–Cdc45 and Sld7–Sld3ΔC from *K. marxianus* (same family as *S. cerevisiae*).

### Supplementary Figure 15

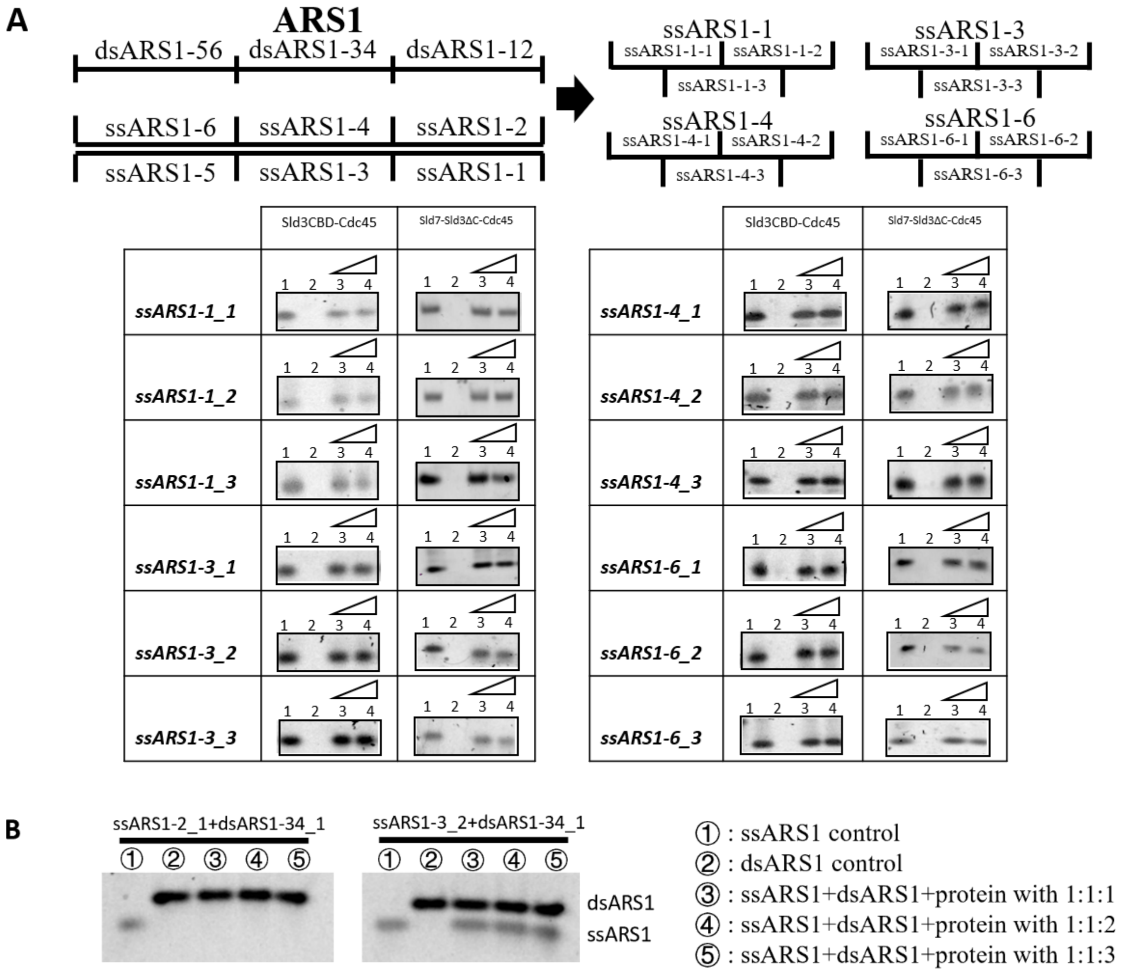

**Supplementary Figure 15. DNA-binding assay by electrophoresis mobility shift assay**

(A) Different concentrations of Sld3CBD–Cdc45 and Sld7–Sld3ΔC–Cdc45 were incubated with ssDNA fragments. Lanes 1, 2, 3, and 4 represent ssDNA to protein ratios of 1:0, 0:1, 1:1, and 1:2, respectively. Lanes 1 and 2 are controls for ssDNA and protein, respectively. The DNAs were visualized using SYBR Safe on polyacrylamide gels. (B) Different concentrations of Sld7–Sld3ΔC–Cdc45 were incubated with the dsDNA fragment (dsARS1-34_1) mixed with jointed ssDNA fragments (ssARS1-2_1 or ssARS1-3_2). The ssARS1-2_1 and ssARS1-3_2 connect to dsARS1-34_1 at different sites. The control for ssDNA and dsDNA is located in lanes 1 and 2, respectively.

**Supplementary Figure 16**

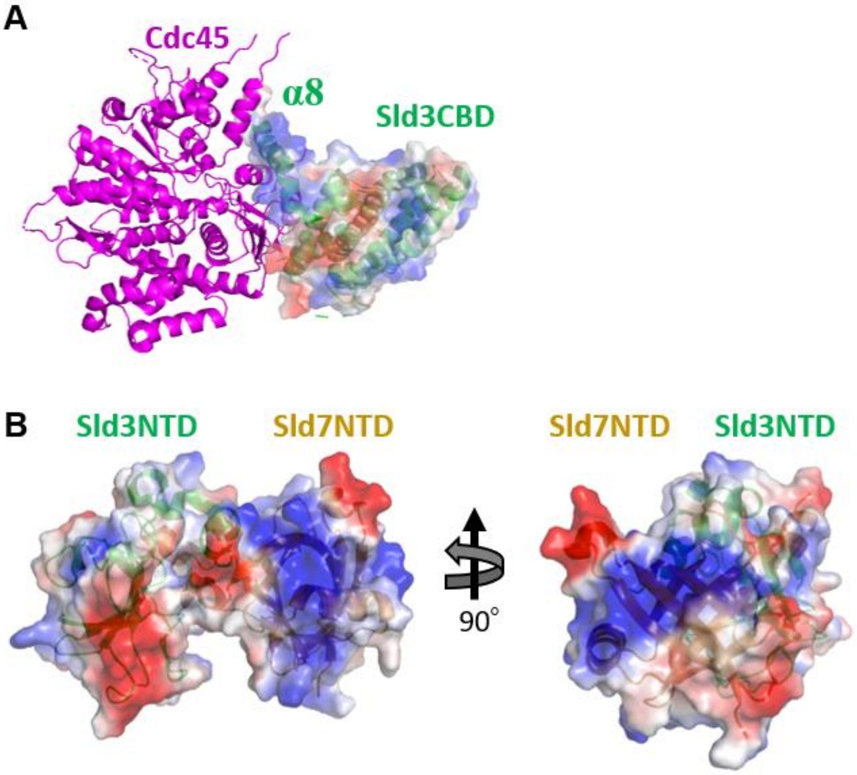

**Supplementary Figure 15. Surface charge of Sld3CBD-Cdc45 and Sld3NTD-Sld7NTD**

The Sld3CBD-Cdc45 (A) and Sld3NTD-Sld7NTD (B) are presented in charged surface calculated by the *Pymol* program. The blue and red show positive and negative charges, respectively. Cdc45 covers the main positive charge area of Sld3CBD α8CTP (A). A large positive charged region surrounds the middle of Sld7NTD in Sld3NTD-Sld7NTD (B).

**Supplementary Figure 17**

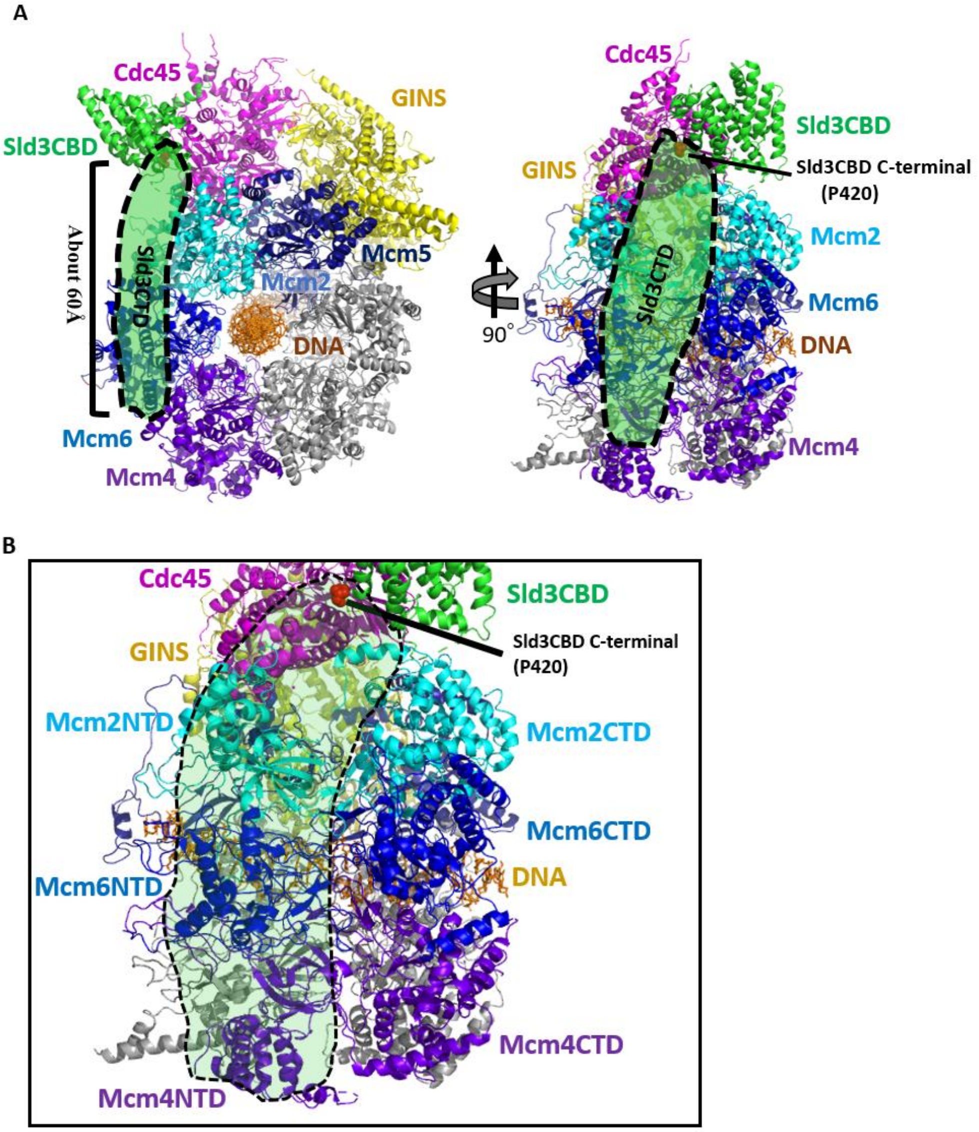

**Supplementary Figure 16. Model of Sld3CTD on the CMG**

(A) Sld3CTD diagram on the SCM complex model. Sld3 could extend the C-terminal domain to interact with the Mcm4 NTD via the NTDs of Mcm2 and Mcm6. (B) Expansion of the Sld3CTD in close-up. The black dotted region denotes the extended region of Sld3CTD, which binds to the NTDs of Mcm2, Mcm4 and Mcm6. Sld3CBD C-terminal P420 is shown as red spheres. Labelled Mcm2, 4, and 6 subunits are colored cyan, blue, and purple blue, respectively. The subunit Mcm5 is colored dark blue, and the subunits Mcm3 and Mcm7 are colored gray. Green and pink are used to color Sld3CBD and Cdc45, respectively. GINS is shown in yellow, and a dsDNA is presented by the stick with dark orange.

**Supplementary Table 1. Primers used in this study**

Sld3CBD-Cdc45

| | **Forward primer** | **Reverse primer** |
|---|---|---|
| **Sld3CBD** | 5'CTCGAGCACCACCACCACCACCACTGAG | 5'GGTATATCTCCTTCTTAAAGTTAAAGTTAAACAAAATTATTTCTAGAGG |
| **Cdc45** | 5'CTCGAGTCTGGTAAAGAAACCGCTG | 5'ATTTCGATTATGCGGCCGTGTACAATAC |

Sld7-Sld3ΔC-Cdc45

| | **Forward primer** | **Reverse primer** |
|---|---|---|
| **Sld7** | 5'GGAATTCCATATGCCGCTGTTTAAAGAAC | 5'CCGCTCGAGTTACGTTTTGGTGAACATTTC |
| **Sld3ΔC** | 5'CATGCCATGGAACCGAGCGAAG | 5'CCGCTCGAGGTTACCCTTCGGAACTTTG |
| **Cdc45** | 5'GGAATTCCATATGTATGGGTATCAACG | 5'CCGCTCGAGTTAAATCAGACCGCTCAGG |

Sld7-Sld3ΔC-Cdc45 IIS

| | **Forward primer** | **Reverse primer** |
|---|---|---|
| **Cdc45 IIS** | 5'ATTATCGAAATCCGCAAAGAAGATTCGCAGCCGTTCTCGGAACGTCTGACCC | 5'GGGTCAGACGTTCCGAGAACGGCTGCGAATCTTCTTTGCGGATTTCGATAAT |

Sld3CBD mutants

| | Forward primer | Reverse primer |
|---|---|---|
| **Sld3-3S** | 5’GACAGTAGTCTATCAAGTGAAAC GCCCAAC | 5’CAAGCTGCATGCTCTATCTAAGTACAAGTC TAACTGTTC |
| **Sld3-3E** | 5’GACGAAGAACTATCAAGTGAAAC GCCCAAC | 5’CAATTCGCATGCTCTATCTAAGTACAAGTC TAACTGTTC |
| **Sld3-Y** | 5’CCCAACCCAGATGCCATAGAAGC AT | 5’CGTTTCACTTGATAGTAGAATGTCCAAGTA GCATGCTCTATC |
| **Sld3-2R** | 5’ACTTACGTAGAGCATGCATCTTG GACATTCTACTATCAAGTG | 5’ACAAGCGTAACTGTTCACAATAATCCAAAG ATGTTGTTATTCTC |

Cdc45 mutants

| | Forward primer | Reverse primer |
|---|---|---|
| **Cdc45-RA** | 5’GCAATTTATAGATTATGCGTCTT ACAAGACGGACCC | 5’TAAATGCTTGATTAATTTCTTCTCCAATATAGCA ACCC |
| **Cdc45-3E** | 5’ATAGAGAGTGCGAGTTACAAG ACGGACCCGATTTAGACTTG | 5’AAATTCTCTCATGCTTGATTAATTTCTTCTCCAA TATAGCAACCC |
| **Cdc45-2E** | 5’CATTCGAAGAGAAGCTGACCT TGAGTGGATTG | 5’GTGATTCATCTTCACGACGTATTTCAATTATGGA ACTTTC |
| **Cdc45-3S** | 5’ATAGAAGTTGCTCATTACAAGA CGGACCCGATTTAGACTTG | 5’AAATTCTTGAATGCTTGATTAATTTCTTCTCCAA TATAGCAACCC |
| **Cdc45-2S** | 5’CATTCAGTGAGAAGCTGACCT TGAGTGGATTG | 5’GTGAACTATCTTCACGACGTATTTCAATTATGGA ACTTTC |
| **Cdc45 W481R** | 5’GCTCAGTAGAGATGCTCTAGAT GACAGAAAGGTGG | 5’ GAGCATCTCTACTGAGCCAAAAATTCGAAACC |
| **Cdc45 G367D** | 5’GCTAGAATGGATATACCATTAA GTACTGCACAAGAAACATG | 5’GGTATatcCATTCTAGCAAACATCTTATGCAATC |

**Supplementary Table 2. Statistics of data collection and refinement**

Statistics for the highest-resolution shell are shown in parentheses

| | |
|---|---|
| Data collection | |
| Resolution range(Å) | 43.5 - 2.6 (2.7 - 2.6) |
| Cell dimension(Å) | 70.8 107.6 128.3 |
| Space group | $P2_12_12_1$ |
| Number of unique reflections | 30622 (3000) |
| Completeness (%) | 99.9 (99.6) |
| Multiplicity | 6.6 (6.5) |
| $R_{merge}$ (%)[a] | 12.7 (97.1) |
| <I/sigma(I)> | 13.3 (2.05) |
| $CC_{1/2}$(%) | 99.7 (75.7) |
| Number of Sld3CBD-Cdc45 in ASU | 1 |
| Wilson B factor | 43.9 |
| Refinement | |
| Resolution range(Å) | 43.5 - 2.6 |
| $R_{free}/R_{work}$ (%)[b] | 26.2 / 21.9 |
| Total number of atoms | 6516 |
| Protein atoms | 6476 |
| Water atoms | 40 |
| Others | 0 |
| Averaged B factor | 55.0 |
| RMS deviations | |
| Bonds (Å) | 0.0026 |
| Angles (°) | 0.63 |
| Ramachandran plot (%) | |
| Favored | 94.1 |
| Allowed | 4.62 |
| Outliers | 1.28 |

[a] $R_{merge} = \sum_{hkl}\sum_i |I_i(hkl) - \langle I_i(hkl)\rangle| / \sum_{hkl}\sum_i I_i(hkl)$, where *i* is the number of observations of a given reflection and *I(hkl)* is the average intensity of the *i* observations.

[b] $R = \sum\left||F_0| - |F_c|\right|/\sum|F_0|$. $|F_0|$ and $|F_c|$ are amplitudes of the observed and calculated structure factors, respectively. $R_{work}$is the R value for reflections used in the refinement, whereas $R_{free}$ is the R value for 5% of the reflections, which are selected in thin shells and are not included in the refinement.
